# A comprehensive large scale biomedical knowledge graph for AI powered data driven biomedical research

**DOI:** 10.1101/2023.10.13.562216

**Authors:** Yuan Zhang, Xin Sui, Feng Pan, Kaixian Yu, Keqiao Li, Shubo Tian, Arslan Erdengasileng, Qing Han, Wanjing Wang, Jianan Wang, Jian Wang, Donghu Sun, Henry Chung, Jun Zhou, Eric Zhou, Ben Lee, Peili Zhang, Xing Qiu, Tingting Zhao, Jinfeng Zhang

**Author notes:** Correspondence: Jinfeng Zhang, **Email:**. These authors contributed equally (Yuan Zhang, Xin Sui, Feng Pan, Kaixian Yu).

## Abstract

To address the rapid growth of scientific publications and data in biomedical research, knowledge graphs (KGs) have become a critical tool for integrating large volumes of heterogeneous data to enable efficient information retrieval and automated knowledge discovery (AKD). However, transforming unstructured scientific literature into KGs remains a significant challenge, with previous methods unable to achieve human-level accuracy. In this study, we utilized an information extraction pipeline that won first place in the LitCoin NLP Challenge (2022) to construct a large-scale KG named iKraph using all PubMed abstracts. The extracted information matches human expert annotations and significantly exceeds the content of manually curated public databases. To enhance the KG’s comprehensiveness, we integrated relation data from 40 public databases and relation information inferred from high-throughput genomics data. This KG facilitates rigorous performance evaluation of AKD, which was infeasible in previous studies. We designed an interpretable, probabilistic-based inference method to identify indirect causal relations and applied it to real-time COVID-19 drug repurposing from March 2020 to May 2023. Our method identified 600-1400 candidate drugs per month, with one-third of those discovered in the first two months later supported by clinical trials or PubMed publications. These outcomes are very challenging to attain through alternative approaches that lack a thorough understanding of the existing literature. A cloud-based platform (https://biokde.insilicom.com) was developed for academic users to access this rich structured data and associated tools.

## Introduction

The sheer volume of information produced daily in scientific literature, expressed in natural languages, makes it impractical to manually read all publications, even within relatively narrow research areas. Additionally, advances in high-throughput technologies have led to the creation of enormous quantities of research data, much of which remains underutilized in various databases. This information explosion poses a major challenge for researchers to identify and develop innovative ideas using all the available data. Automated knowledge discovery (AKD, a.k.a. automated hypothesis generation) can help mitigate this problem by automating the process of data analysis, identifying patterns, and generating innovative insights and hypotheses^1^. In recent years, knowledge graphs (KGs) have been proposed as a powerful data structure for integrating heterogeneous data and for AKD^2–8^. KGs, with entities as nodes and their relationships as edges, represent human knowledge in a structured form, facilitating efficient and accurate information retrieval. Graph algorithms can be employed on KGs to infer potential relationships as plausible hypotheses between known entities.

Computational construction of KGs from unstructured text entails two steps: named entity recognition (NER) to identify key biological entities and relation extraction (RE) to extract relationships among entities. Historically, NER and RE have been collectively referred to as information retrieval tasks. Early automated methods mainly fell into two categories: rule-based and machine learning-based. The rule-based approach systematically extracted specific data based on predefined rules^9–14^, while the machine learning-based approaches inferred rules from annotated data usually with increased recall and overall performance^15–30^. The advent of machine learning led to more sophisticated methods that leveraged semantic information and sentence structure, resulting in significant improvements in information extraction effectiveness^21,24^. However, a gap remained compared to human proficiency.

The emergence of deep learning models has allowed for a more nuanced utilization of information, such as semantic content and grammatical structures. By expanding the use of features and enhancing expressive capabilities, deep models have significantly improved the overall effectiveness of information extraction^31–43^. Recently, the technique of pretraining and large language models (LLMs) have garnered considerable attention, expanding both model complexity and the amount of training data and achieving remarkable progress in information retrieval tasks^43–54^. This was evidenced by the significant results in the BioCreative VII Challenge in 2021, where finetuning BERT-based models was widely used, and the top performance in some tasks closely matched human annotator performance. Subsequently, a highly advanced series of pre-trained models, like GPT-4, emerged^55–57^. These models have been proven to perform comparably or better to humans in multiple tasks, marking a significant breakthrough in the field.

Recently, LLMs like GPT-4 have been explored for their integration with KGs, aiming to enhance tasks such as named entity recognition (NER), relation extraction, and event detection through techniques like zero-shot prompting, in-context learning, and multi-turn question answering^58–60^. While these models excel in generalization and large-scale data processing, they still struggle with domain-specific challenges, including handling long-tail entities^61^, directional entailments^62^, and inconsistencies in retrieving knowledge from paraphrased or low-frequency phenomena^63,64^. Experiments^60^ on datasets like DuIE2.0^65^, Re-TACRED^66^, and SciERC^67^ highlight that fine-tuned small models continue to outperform GPT-like LLMs in KG-related tasks. Despite these limitations, LLMs have shown significant adaptability in augmenting KGs, particularly when structured data is limited, positioning them as a complementary tool for AKD^58^.

To facilitate the methodology development and identification of the most effective methods for KG construction, the National Center for Advancing Translational Sciences (NCATS) of the National Institutes of Health (NIH) organized the LitCoin natural language processing (NLP) challenge between Nov 2021 and Feb 2022. In the LitCoin NLP Challenge dataset, six common biological entity types were annotated: diseases, genes/proteins, chemical compounds, species, genetic variants, and cell lines. Eight relation types were also annotated for the entities: association, binding, comparison, conversion, cotreatment, drug interaction, positive correlation, and negative correlation. These entity types and relations are highly relevant in translational research and drug discoveries. Our team, JZhangLab@FSU, participated in the challenge and won first place (https://ncats.nih.gov/funding/challenges/winners/litcoin-nlp).

In this study, we applied our LitCoin NLP Challenge-winning information retrieval pipeline to all PubMed abstracts (cutoff date: May 2023) to construct a large-scale Biomedical Knowledge Graph (iKraph, the abbreviation for Insilicom’s Knowledge Graph). Manual verification confirmed the pipeline’s accuracy at a human annotator level. By annotating the directions for the relations in the LitCoin dataset and training a model to predict the direction of relations, we constructed a causal knowledge graph (CKG) capable of making indirect causal inferences. To further enhance the coverage of iKraph, we integrated relation data from public databases and high-throughput genomics datasets, making it the most comprehensive, high-quality biomedical knowledge graph to date. To make causal inferences among the entities that are not directly connected in the KG, we designed a probabilistic-based approach, probabilistic semantic reasoning (PSR). PSR is highly interpretable as it directly infers indirect relations using direct relations through straightforward reasoning principles.

Navigating the modern drug development terrain is intricate and resource-intensive^68^. The ascent in costs largely stems from prior research exhausting more straightforward drug targets, necessitating a shift towards more complex ones^69^. In this setting, knowledge graphs play a pivotal role in automated knowledge discovery (AKD)^2,70–72^, particularly in the domain of drug target identification and drug repurposing^73–77^. A significant challenge in developing methods for such applications has been to comprehensively assess the effectiveness of these studies. For example, in the case of drug repurposing, collecting all the known therapeutic associations of a particular disease or drug requires a thorough search of the literature. Without such knowledge, it is impossible to rigorously evaluate drug repurposing methods. In our investigation, for each repurposing objective, we extracted all therapeutic associations documented in PubMed abstracts. This enabled us to measure recall and observed positive rate (OPR), which is infeasible in prior drug repurposing research.

We demonstrate the power of our approach by conducting several drug repurposing studies: drug repurposing for COVID-19, cystic fibrosis, ten diseases without satisfactory treatment, and ten commonly prescribed drugs. For COVID-19 and cystic fibrosis, we performed retrospective, real-time drug repurposing exercises. Our method identified numerous viable candidates, supported by substantial literature evidence connecting the drug and disease entities. This level of interpretability is invaluable when determining the necessity of subsequent research endeavors.

## Results

### Building a Large-Scale Biomedical Knowledge Graph (iKraph)

To facilitate the methodology development and identification of the most effective methods for KG construction, NIH organized the LitCoin natural language processing (NLP) challenge between Nov 2021 and Feb 2022 (https://ncats.nih.gov/funding/challenges/litcoin). Our team, JZhangLab@FSU, participated in the challenge and won first place (Table 1). In the summer of 2023, we also participated in the BioRED track of the BioCreative Challenge VIII. In the end-to-end KG construction task, our team also achieved the highest score (Table 1)^78^. The LitCoin NLP Challenge dataset comprises 500 PubMed abstracts, each annotated with six distinct entity types and eight types of relations at the abstract level. We used the pipeline initially developed for the LitCoin challenge to process all PubMed abstracts available before May 2023, creating a large-scale Knowledge Graph, iKraph. In constructing iKraph, we processed over 34 million PubMed abstracts, resulting in 10,686,927 unique entities and 30,758,640 unique relations. We incorporated entity normalization into our pipeline, as this was not a component of the LitCoin NLP challenge (see Supplementary Materials Section 1.2, 1.3 for more details).

**Table 1:** The Performance of the top teams in the LitCoin NLP challenge and BioCreative challenge VIII (BC8) BioRED track subtask 2(End-to-End KG construction). The LitCoin NLP Challenge results include NER and RE scores, reported as Jaccard scores, and the overall score. Our team, JZhangLab@FSU, achieved the highest overall score. The BioCreative VIII BioRED track Subtask 2 results present F1 scores for various evaluation metrics. Our team, Insilicom, ranked first. ‘+’ indicates additional task-specific scores. Bold values indicate the highest-performing scores within each column in each competition.

| Competition | Team | NER | RE | Overall | +ID | +Entity pair | +Relation type | +Novelty |
| --- | --- | --- | --- | --- | --- | --- | --- | --- |
| LitCoin | JZhangLab@FSU | 0.9177 | **0.6332** | **0.7186** | - | - | - | - |
|  | UTHealth SBMI | **0.9309** | 0.5941 | 0.6951 | - | - | - | - |
|  | UIUCBioNLP | 0.9068 | 0.5681 | 0.5681 | - | - | - | - |
| BC8 | 156 (Insilicom) | **0.8926** | - | - | 0.8407 | **0.5584** | **0.4303** | **0.3275** |
|  | 129 | 0.7858 | - | - | 0.7635 | 0.4127 | 0.3103 | 0.2334 |
|  | 127 | 0.7830 | - | - | 0.7598 | 0.3945 | 0.2976 | 0.2280 |
|  | 118 | 0.7831 | - | - | 0.7561 | 0.3903 | 0.2859 | 0.2248 |
|  | GPT-3.5 + PubTator3 | 0.8652 | - | - | **0.8565** | 0.3021 | 0.1081 | 0.0714 |
|  | GPT-4+PubTator3 | 0.8652 | - | - | **0.8565** | 0.3449 | 0.1704 | 0.1277 |

We evaluated the accuracy of our large-scale relation extraction (RE) and our novelty prediction results using a sample of 50 randomly selected PubMed abstracts, including 1583 entity pairs. The results shown in Supplementary Table 3 indicate that our information extraction performance rivals that of human annotations. A more in-depth analysis is available in the Supplementary Materials Section 2.

Fig. 1A shows the number of PubMed abstracts containing one or more of the four major types of entities: diseases, genes, chemicals, and sequence variants. It is evident that diseases are the most common topic, with over 20 million articles referencing at least one disease entity, and nearly half of these articles focus exclusively on diseases. In contrast, gene mentions often coexist with other entities, such as chemicals and diseases. Fig. 1B depicts the number of PubMed abstracts containing one or more of the five major types of relations, offering insight into the distribution of topics in biomedical research.

**Fig. 1.**
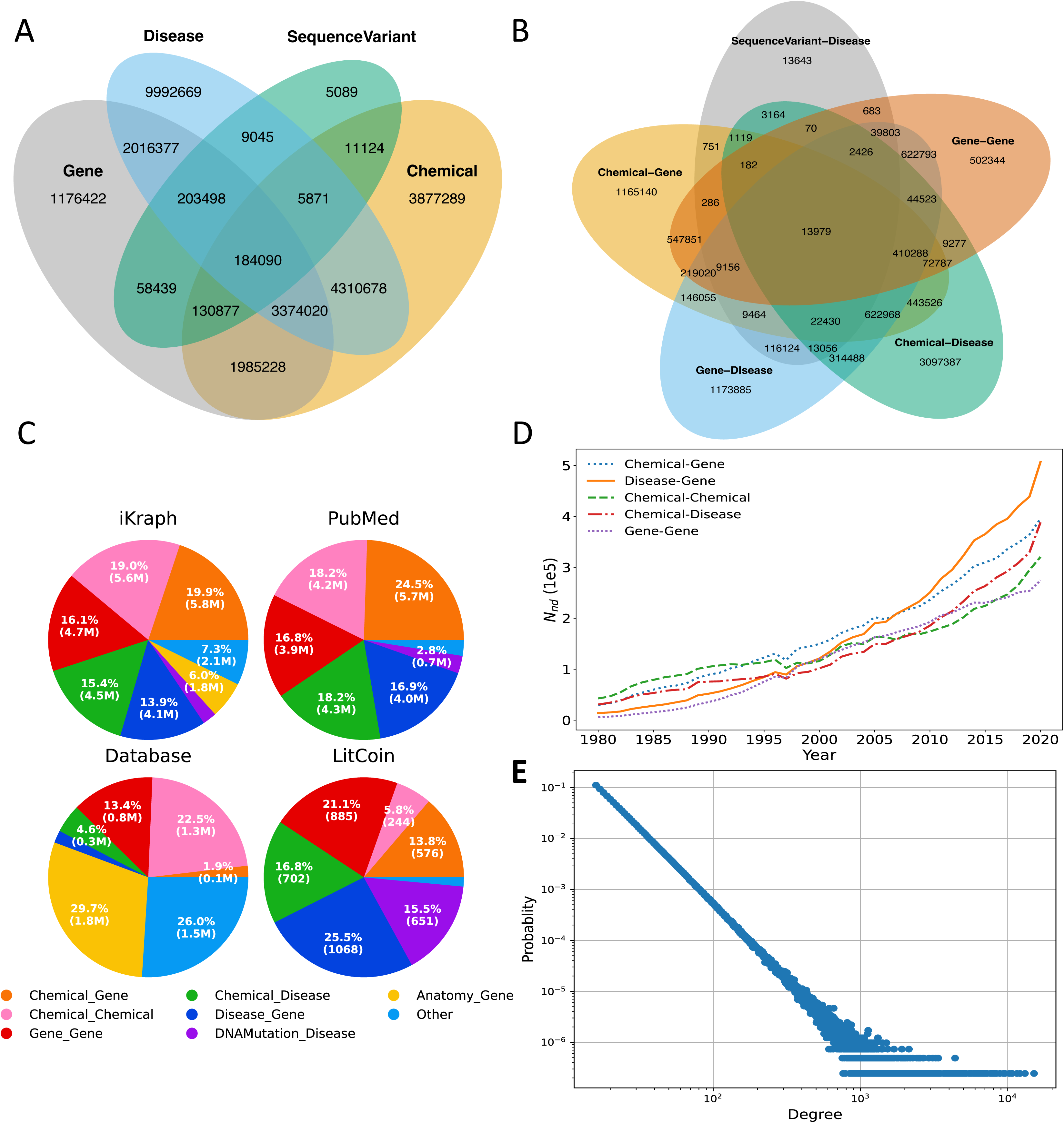
**A**. Venn diagram for the number of PubMed articles containing certain types of entities; B. Venn diagram for the number of PubMed articles containing certain types of relations; **C**. The composition of relations in iKraph, PubMed abstracts, public databases, and LitCoin dataset; **D**. The numbers of novel discoveries by entity pair type from 1980 to 2020. **E**. Degree distribution of iKraph, where the x-axis represents the degree k of an individual entity, and the y-axis p(k) denotes the corresponding probability of any entity exhibiting that degree.

Fig. 1C compares the relations extracted from PubMed with those from databases and the LitCoin dataset. There is a clear difference between the LitCoin dataset and general PubMed abstracts, as the former contains more relations in each abstract, especially those involving sequence variant entities^79^. This explains the performance difference of our pipeline on these two datasets. Relations from PubMed and public databases are also quite complimentary.

Fig. 1D shows the number of novel discoveries for different entity pairs over the year. We have observed a remarkable upswing in disease-gene relations since 2005, which underscores the tangible outcomes of translational initiatives promoted by federal agencies. Furthermore, the increasing number of disease-gene relations signifies an improved understanding of disease mechanisms at the molecular level, thereby bolstering efforts in drug discovery. Of particular note is the rapid escalation of chemical-disease relations in recent years, particularly around 2020, which is anticipated to continue in the foreseeable future.

We plotted *p*(*k*) vs *k*, where *k* is the degree of an entity in iKraph, and *p*(*k*) is the probability of an entity having degree *k* (Fig. 1E). We found that iKraph exhibits a scale-free topology with an alpha parameter value of around 3.0 (more details in Supplementary Materials Section 3).

Supplementary Table 4 compares the numbers of relations for five types of entity pairs from all the public databases integrated into iKraph, those extracted from PubMed, and the numbers extracted if we use a simple co-occurrence rule, which considers two entities having a relation if they co-occur in an abstract. On the one hand, iKraph has significantly more numbers of relations than those from public databases. On the other hand, the numbers of co-occurrences are much larger than relations extracted from PubMed, indicating a substantial noise reduction by explicitly extracting relations from literature compared to retrieval using keywords.

### Constructing a causal knowledge graph

We developed a model to predict the direction of correlation relations in the LitCoin dataset, identifying whether the relation flows from entity1 to entity2 or entity2 to entity1. Adding this directional information transformed correlations into potential causal relationships, allowing us to construct a directed knowledge graph for knowledge discovery applications.

### PSR for inferring indirect relationships

With directional information, we can infer relations between indirectly connected entities using straightforward reasoning. To this end, we designed the probabilistic semantic reasoning (PSR) algorithm, which is both efficient and interpretable. PSR enables all-against-all drug repurposing for all drugs and diseases with limited computational resources and allows efficient updates of newly inferred relations.

For instance, freshly published PubMed articles can be processed daily to extract discoveries and generate hypotheses for timely dissemination. In contrast, most machine learning methods struggle to achieve this level of efficiency and interpretability.

### Drug repurposing for Covid-19 using iKraph

Using the PSR algorithm, we conducted a retrospective, real-time drug repurposing study for COVID-19 spanning from March 2020 to May 2023 (Fig. 2). During this period, we consistently discovered repurposed drugs based on the drug targets reported for COVID-19 between March and June 2020. A candidate drug has at least one directed path to COVID-19 through an intermediate gene. We checked whether any repurposed drugs were later validated by either PubMed literature or clinical trials through a monthly assessment. The validation involved scrutinizing whether these repurposed drugs had been subsequently tested in clinical trials documented on ClinicalTrials.gov or had published therapeutic efficacy in COVID-19 patients in PubMed abstracts. Notably, drugs identified in clinical trials may not always translate into effective treatments for COVID-19. Nevertheless, they serve as valuable hypotheses, aligning with the primary objective of our drug repurposing approach. As shown in **Error! Reference source not found.**A, we were able to identify nearly 600 to 1,400 candidate drugs from iKraph using PSR. Remarkably, one-third of the repurposed drugs identified during the initial two months were later validated as effective treatments or plausible potential treatments worthy of clinical trials. Importantly, even drugs that did not achieve validation status remain viable hypotheses, warranting further investigation, particularly when existing treatments prove less than optimal.

**Fig. 2.**
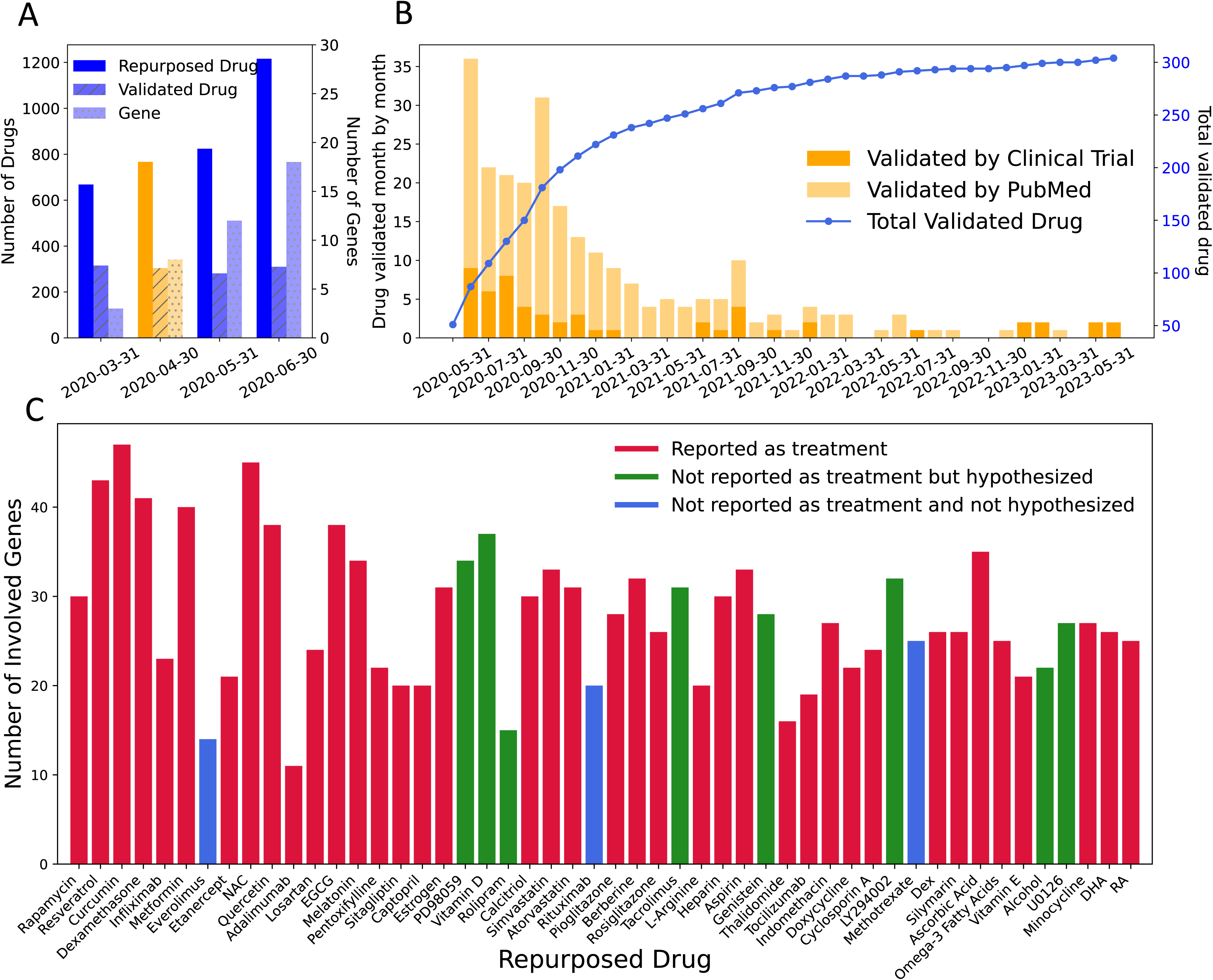
Drug repurposing for COVID-19. **A.** The number of repurposed drugs, number of verified drugs, and number of COVID-19-related genes for the first four months of the COVID-19 pandemic (March to June 2020); **B.** The number of verified drugs each month for those repurposed for Apr. 2020; **C.** The number of genes involved in the drugs repurposed at present time (March 2023). The figure shows the top 50 repurposed drugs sorted from left to right, with those on the left having higher scores. Almost all the repurposed drugs interact with many genes (height of the bar) related to COVID-19. The majority of the drugs were reported as treatment for COVID-19 (36 out of 50). Among those that were not reported as treatments for COVID-19, 11 out of 14 were hypothesized as potential treatments.

Fig. 2B shows a timeline of repurposed drug validation. Notably, there is a surge in validated drugs during the first year, which subsequently shows a month-to-month decline. This trend suggests most repurposed drugs align with practitioners’ early assessments. Some drugs were validated only in the second or third year, indicating they were less immediately evident. The number of drugs validated through publications matches those validated via clinical trials. While numerous drug repurposing studies for COVID-19 exist^80–83^, as per our understanding, no prior research has as thoroughly validated such a vast quantity of repurposed drugs as we have in this research. These results highlight iKraph’s ability to identify promising drug candidates for specific diseases in real-time.

We then conducted drug repurposing for COVID-19 in the current timeframe (Fig. 2C). We did not exclude drugs already reported as treatments for COVID-19 (direct relations). This was to observe if our repurposing efforts agree with existing treatment choices for COVID-19. Fig. 2C displays the top 50 repurposed drugs. Notably, most of these drugs (36 out of 50) have published studies mentioning either their potential therapeutic efficacy or demonstrated therapeutic efficacy for phenotypes associated with COVID-19. Among the remaining 14, 11 have been proposed as potential treatments for COVID-19 (citations provided in Supplementary Table 3). For each drug, numerous genes that link COVID-19 with the drug were identified (y-axis of Fig. 2C). Additionally, each of these relations, whether drug-gene or gene-COVID-19, is supported by one or multiple articles. To our knowledge, none of the previous literature-based COVID-19 repurposing studies has yielded such comprehensive findings.

### Drug repurposing for cystic fibrosis using iKraph

We applied PSR to uncover indirect relationships between drugs and cystic fibrosis (CF) from 1985 to 2022 (Fig. 3). Since the early 1990s, at least 50 potential repurposed drugs have been identified annually. A drug was considered validated if later reported as directly therapeutic for CF. Historically, estimating these metrics was challenging due to reliance on manual literature searches. We calculated recall (percentage of known direct relations successfully repurposed) and observed positive rate (OPR, percentage of repurposed cases with reported direct relations). Unlike precision, OPR accounts for potential candidates awaiting validation. From 1990 to 2022, the average recall is 0.635 (Fig. 3B), and the average OPR from 1985 to 2011 is 0.159. Different time intervals were used because OPR requires earlier predictions, while recent predictions need time for validation.

**Fig. 3.**
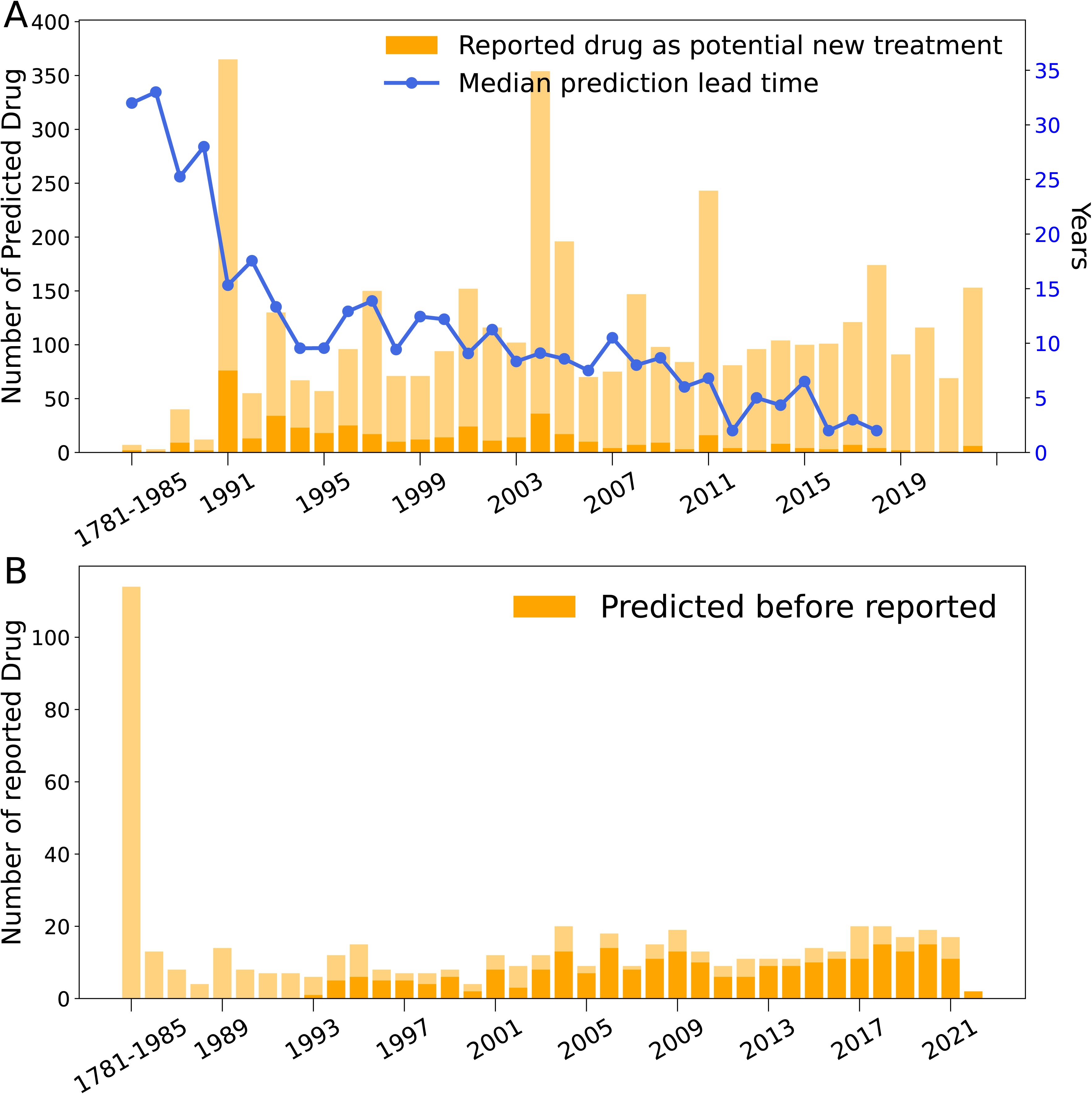
Drug repurposing for cystic fibrosis. **A.** The number of repurposed drugs from 1985 to 2022. The dark yellow bar shows the number of validated drugs. The blue line shows the time it takes to validate the targets for the corresponding years. **B.** The number of drugs reported in PubMed from 1985 to 2022. The dark yellow bar shows the drugs that have been predicted previously.

We calculated the typical duration for these repurposed drugs to be validated. Remarkably, our proposed drugs typically appeared in literature 2 to 33 years later, with a median validation time of 9.4 years (Fig. 3A). Assuming experimental validation takes 2 years on average, iKraph could hypothetically reduce this time from 9 to 2 years if predictions were acted on immediately. With over 63% recall and a 9-year median lag, our findings highlight iKraph’s potential to accelerate drug repurposing and validation for cystic fibrosis treatment.

### Drug repurposing for 10 diseases and 10 drugs

To evaluate our method’s versatility, we extended drug repurposing to ten diseases lacking satisfactory treatments and ten commonly prescribed drugs (Supplementary Figure 3). Our PSR algorithm identified a vast array of candidates for these drugs and indications. For each drug (or disease) assessed, we calculated both the recall and the observed positive rate (OPR). Impressively, our findings revealed average recall values of 0.76 for disease repurposing and 0.86 for drug repurposing. This exceptional recall rate emphasizes the potency of iKraph coupled with our PSR algorithm in spotlighting viable drug repurposing candidates. Notably, these elevated recall rates were achieved without an excessive number of predictions. The observed OPRs remained commendable at 0.197 for diseases and 0.07147 for drugs.

Importantly, a significant proportion of indications repurposed for these drugs are not associated with any treatments in PubMed abstracts. This suggests that certain ailments might still be without treatments, and these widely used drugs could potentially fill those therapeutic gaps.

We extended our analysis by using relations from a database, alongside those extracted from PubMed abstracts, to make drug repurposing predictions. Fig. 4 illustrates the F1 scores for these comparisons based on the top 50 predicted repurposed drugs and the top 250 predicted indications. In each panel, blue bars represent PubMed-based predictions, while orange bars represent database-based predictions. Most repurposed drugs or diseases showed higher F1 scores using PubMed-derived predictions, likely due to the greater amount of information available in PubMed, which databases cannot match.

**Fig. 4.**
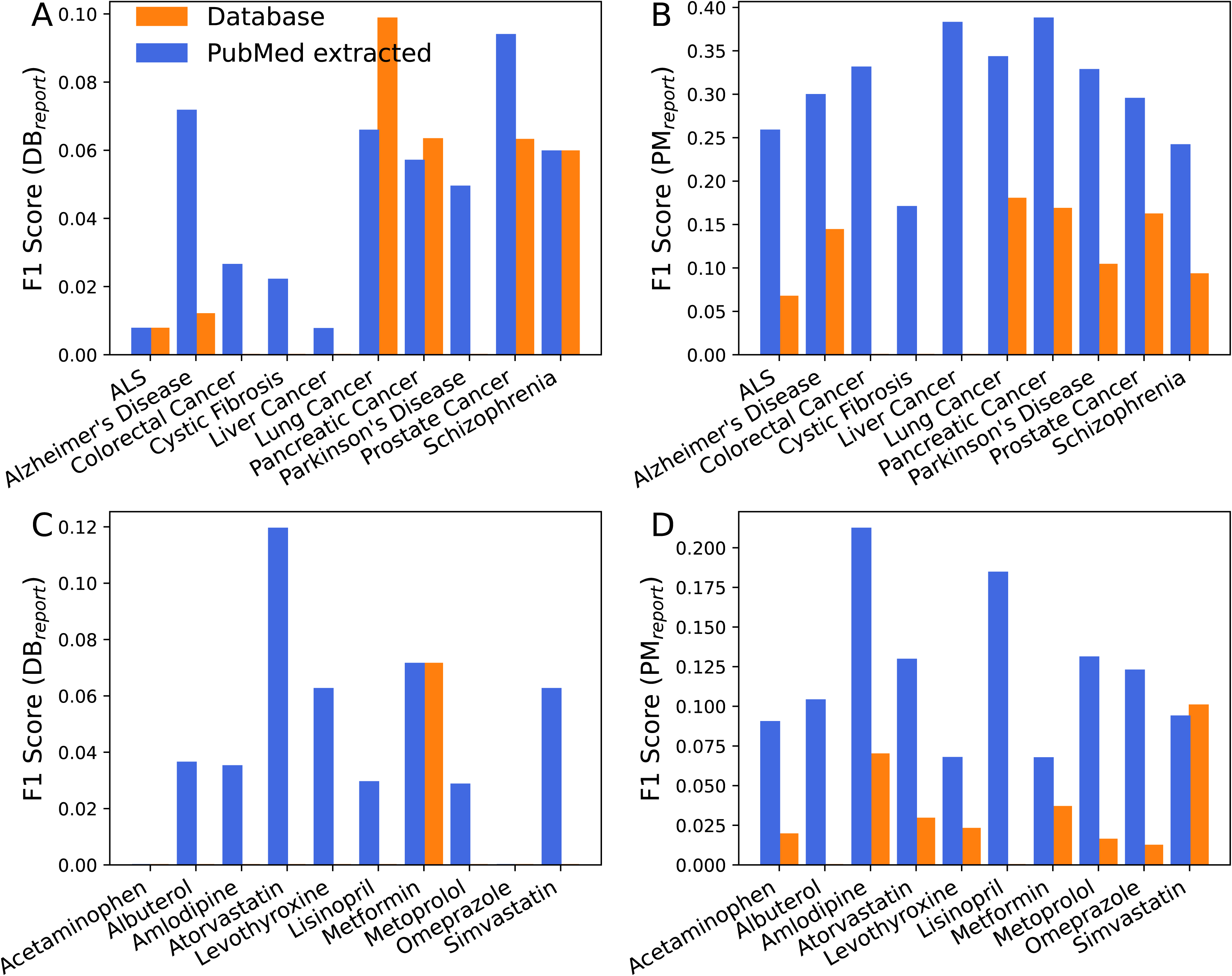
F1 scores for drug repurposing prediction for 10 diseases and 10 common drugs. The calculation is based on the top 50 repurposed drugs and the top 250 repurposed indications. The blue bars represent predictions made using relations extracted from PubMed abstracts, and the orange bars represent predictions made using relations from the database. Panels A and C are validated using therapeutic relations reported in the database, while panels B and D are validated using therapeutic relations extracted from PubMed abstracts.

## Discussion

Converting unstructured scientific literature into structured data has been a long-standing challenge in natural language processing (NLP). Successfully addressing this issue can potentially revolutionize the pace of scientific discoveries. Although numerous studies have been conducted over the years, computational methods have yet to achieve the precision of manual annotation in relation to extraction, posing a significant hurdle. The emergence of LLMs in recent years has ushered in noteworthy advancements in information extraction through LLM fine-tuning. In this paper, we report the first utilization of a human-level information extraction pipeline to construct a large-scale biomedical knowledge graph by processing all the abstracts in PubMed. By further integrating relation data from 40 public databases and those analyzed from publicly available genomics data, the resulting knowledge graph, dubbed iKraph, stands out as perhaps the most all-encompassing biomedical knowledge graph constructed so far. The coverage of iKraph is much larger than public databases for the relations we have extracted. The construction of a causal knowledge graph and the design of an interpretable PSR algorithm allows us to perform automated knowledge discovery very effectively. The exhaustive nature of iKraph allows us to perform research that was infeasible previously. For the first time, we were able to evaluate the performance of automated knowledge discoveries systematically and rigorously by calculating recalls and observed positive rates (OPRs). Without the knowledge of all PubMed abstracts in a structured form, one must perform a manual search of the literature, which would not be feasible for a relatively large number of predictions. We summarize the notable advances in this study, including some unique iKraph-enabled capabilities in Supplementary Material Box S1, and discuss some of them below.

The biomedical research community has traditionally invested significant resources and human effort in knowledge curation through manual annotations. Our research suggests a paradigm shift, leveraging the capabilities of modern LLMs. By initially producing a limited set of high-caliber labeled data, it is feasible to train an information extraction model that operates at human-level precision on much larger text datasets. This methodology could notably expand the reach of public databases without compromising data quality.

Utilizing iKraph for knowledge discovery tasks, such as drug repurposing, has yielded a vast array of credible candidates supported by an unparalleled volume of literature evidence. This underscores the potential of structured knowledge in hastening scientific breakthroughs. In our drug repurposing endeavors for COVID-19, we highlighted iKraph’s proficiency in identifying treatments for pandemics, marking it as an indispensable asset for potential future outbreaks.

Many users might inquire about how iKraph handles noisy information from low-quality papers. Our approach involves aggregating the probabilities of relations (e.g., between A and B) across multiple papers. Each paper assigns a probability to a specific relation, and these probabilities are combined to form an overall score. The more papers that mention the relation, the higher the final probability, making it less susceptible to noise. Relations with low-quality evidence tend to appear in fewer papers, resulting in lower scores. However, while this method provides a strong foundation for handling noisy data, future improvements could involve weighting papers based on factors like journal impact factor, citation count, and publication date. Integrating such metrics aligns with approaches demonstrated in prior work, where features like author diversity, institutional independence, and publication density were found to predict the robustness and reproducibility of scientific claims^84^. Integrating these metrics would allow us to refine the score further by giving more weight to higher-quality sources. For example, papers published in high-impact journals or those widely cited in the scientific community may provide stronger evidence for a relation than those from less reputable sources. Additionally, the publication date can be factored in to balance the relevance of older versus newer findings, ensuring that the most current and impactful research plays a more prominent role in shaping the final probability. This reflects insights from robust scientific literature, where combining high-throughput experimental data with features predictive of reliability has shown promise in assessing the reproducibility of claims^84^. This holistic approach would help iKraph remain robust against misinformation while continuously improving the accuracy of its predictions through adaptive weighting.

Finally, we would like to put our study in the context of the LLMs popular in the current NLP research community. While LLMs have showcased exceptional capabilities in understanding and generating natural language, they aren’t without shortcomings. A notable limitation is their fixed knowledge cut-off date, which restricts their awareness of the very latest developments. Furthermore, in biomedical research, where precision is crucial, relying solely on LLMs to answer specific questions risks inaccuracies due to their limited knowledge base. Additionally, LLMs possess a propensity to generate text that, while convincingly articulated, may lack factual accuracy. This propensity raises concerns regarding the veracity of answers generated by LLMs, necessitating mechanisms for verification and the production of more substantiated results, possibly with appropriate citations. We believe that integrating knowledge graphs like iKraph with LLMs can effectively mitigate these limitations. To this end, we are actively developing a comprehensive question-answering system, combining iKraph with an open-source LLM.

In the Supplementary Materials Section 6, we delve into future research avenues and the challenges we’ve faced. In summary, iKraph serves as a powerful enabler for more effective and efficient information retrieval and automated knowledge discovery.

## Methods

### 1. Information extraction pipeline

We utilized the pipeline crafted during the LitCoin NLP Challenge (https://ncats.nih.gov/funding/challenges/litcoin) to process all PubMed abstracts available until 2022, along with data from several renowned biomedical databases, leading to the creation of the Knowledge Graph, iKraph. The construction of iKraph involves three primary stages: named entity recognition (NER), relation extraction (RE), and novelty classification. The details of the methods can be found in the Method section in the Support Information. When developing the pipeline for LitCoin Challenge, we tested a large set of pre-trained language models including BERT^46^, BioBERT^48^, PubMedBERT^85^ abstract only, PubMedBERT fulltext, sentence BERT^86^, RoBERTa^87^, T5^88^, BlueBERT^89^, SciBERT^54^, and ClinicalBERT^90^. We tested many ideas, such as different loss functions, data augmentations, different settings of label smoothing, different ways of ensemble learning, etc. Our final pipelines contain the following components: (1) Improved in-house script for data processing, including sentence split, tokenization, and entity tagging; (2) RoBERTa large and PubMedBERT models as baseline models for NER task; (3) Ensemble modeling strategy that combines models trained with different parameter settings, different random seeds and at different epochs for both NER and RE; (4) Label smoothing for both NER and RE; (5) Using Ab3P^91^ for handling abbreviations for NER; (6) Modified classification rule tailored for LitCoin scoring method; and (7) Training a multi-sentence model for predicting relations at document level, which gave a very competitive baseline for relation extraction.

We used the pipeline developed in the LitCoin challenge to process all the abstracts in the PubMed database, which contains over 34 million abstracts, resulting in 10,686,927 unique entities and 30,758,640 unique relations.

### 2. Constructing a causal knowledge graph

To infer causal relations, we first annotated causal direction for 4,572 relations in the LitCoin dataset. Among them, 2,009 cases have direction from the first entity to the second; 1,611 cases have direction from the second entity to the first; and 952 cases have no direction. This annotation allowed us to train a model for predicting the directions for relations, which achieved an F1 score of 0.924 in a 5-fold cross-validation test on the LitCoin dataset. Using a causal knowledge graph, we can infer indirect causal relations more effectively for entities not directly connected in the knowledge graph – an essential task in automated knowledge discovery.

To make inferences using the causal knowledge graph, we designed a probabilistic semantic reasoning (PSR) algorithm, which calculates the probability of a true relation between two entities connected directly or indirectly. For two entities with a direct edge (a relation mentioned in the literature), there can be multiple mentions in different articles. It is necessary to estimate an overall probability for this pair, which will be used for estimating probabilities for indirectly connected entity pairs. PSR is highly interpretable, which is key for the validation of predictions.

The overall drug repurposing strategy and validation approach are depicted in Fig. 5, with some details provided in the figure legend.

**Fig. 5.**
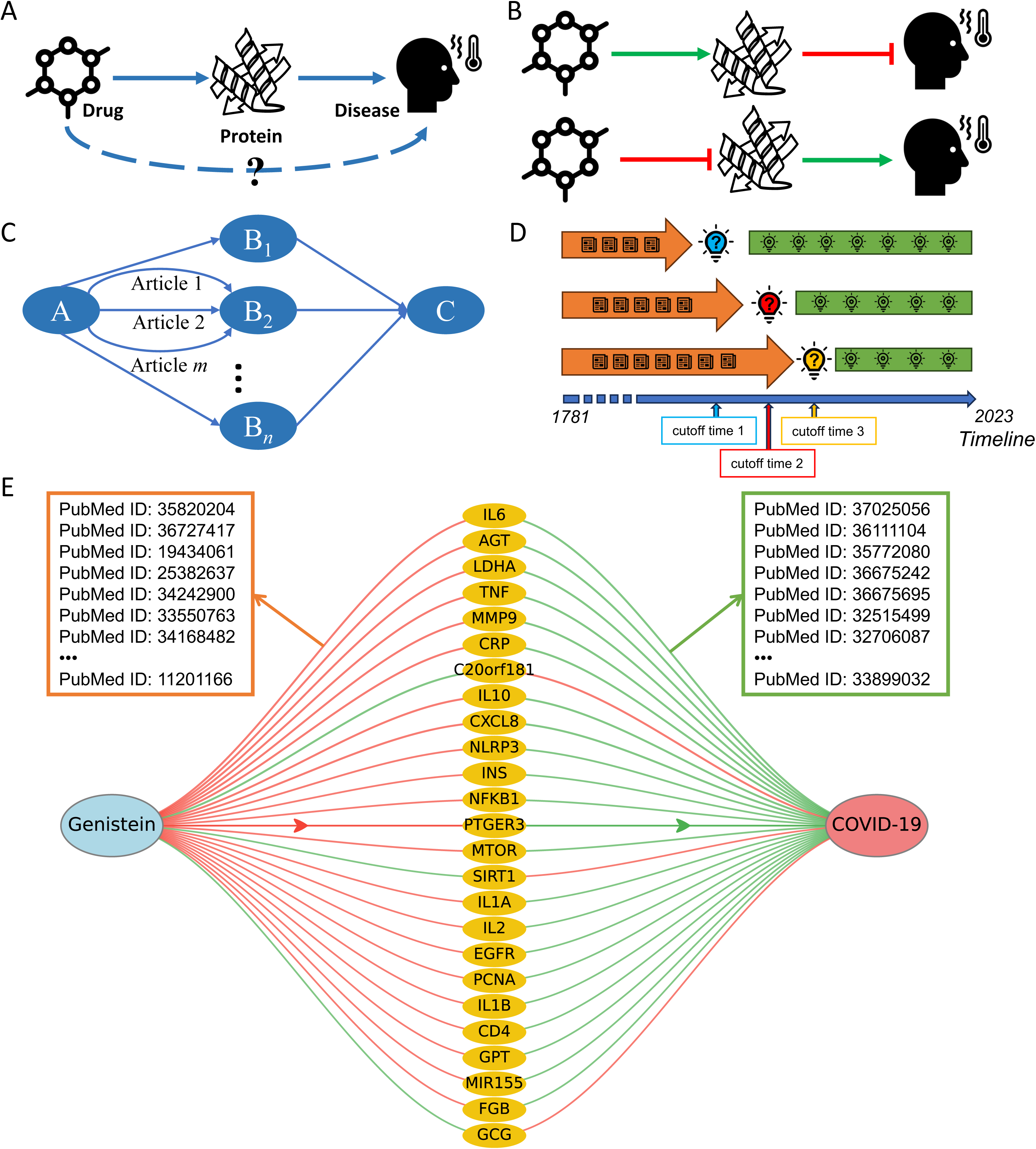
The overview of our drug repurposing strategy and validation approach. **A.** Our method infers drug-disease therapeutic relations through identifying an intermediate entity, the drug target of the disease, with causal relations from the drug to a drug target, and from the drug target to the disease. **B.** Two scenarios correspond to a drug-disease therapeutic relation: drug activates a target, and the target represses the disease, or drug inhibits the target, and the target promotes the disease. **C.** To infer an indirect relation from A (i.e. a drug) to C (i.e. a disease), there can be many intermediate potential targets between them. For each relation formed by two entities, there could be many scientific articles mentioning it. The probabilistic semantic reasoning (PSR) algorithm aggregates all the information to make the inference, while identifying polypharmacological candidates. **D.** To validate a drug repurposing study, we use a time-sensitive approach. We select many cutoff time points (lightbulb with a question mark) and use the knowledge published before the cutoff time (indicated by an orange arrow) to generate predictions and use the knowledge published after the cutoff time (lightbulbs on the green arrow) to validate our predictions. The figure shows three sets of discoveries (blue, red, and yellow bulbs) using three different cutoff times. **E**. Genistein’s repurposing for COVID-19 treatment. Connected via 25 human genes represented by yellow ovals. The diagram shows drug-gene and gene-disease correlations, with red lines indicating negative correlations and green lines positive ones. Each connecting line is supported by multiple articles from which the correlation probabilities are derived.

#### 2.1 Probabilistic Semantic Reasoning (PSR)

To make inferences using the causal knowledge graph, we designed a probabilistic framework, probabilistic semantic reasoning (PSR), for inferring indirect causal relations. PSR is highly interpretable, which is critical for the validation of predictions. There can be multiple mentions in different articles of two entities with a direct edge (a relation mentioned in the literature). It is necessary to estimate an overall probability for this pair, which will be used for estimating probabilities for indirectly connected entity pairs.

To simplify the discussion, let’s assume we want to infer the indirect relation from A to C using direct relations from A to B and the relation from B to C. To infer the indirect relation, we first extract the two direct relations. As mentioned earlier, relation A to B and B to C will likely occur many times in different PubMed abstracts. We calculate the overall probability of whether two entities have a particular relation using the formula:

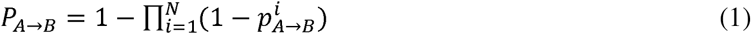

In Equation (1), *P_A→B_* is the overall probability of A-B entity pair having a particular relation, 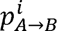 is the probability of being true for the i-th occurrence of these two entities in a PubMed abstract, 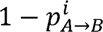 is the probability of this occurrence being false, and 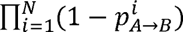is the probability that all the occurrences being false (assuming the predictions for these occurrences are independent). The probability of all occurrences being false, when subtracted by 1, gives the probability that at least one of them is true, which is the desired probability. It is also possible that several different relation types will be inferred for a single pair of entities. Often, only one relation type is true, and others may be wrong predictions. To simplify the inference, we selected the relation type with the highest probability as the true relation type for any pair of entities. In reality, there can be multiple entities linking A to C. We denote one of them as B_j_. Then, the probability of A to C through B_j_ can be calculated as:

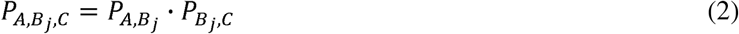

Equation (2) is straightforward since for the indirect relation between A and C (direction from A to C) to be true, both the direct relations must be true. Again, we assume the predictions for the two direct relations are independent. The probability between A to C through m intermediate nodes can then be calculated as

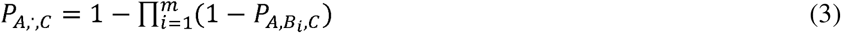

In Equation (3), we use P_A,·,C_ to denote the probability that the indirect relation between A and C through any intermediate entity and there is m such intermediate entities that link A and C. The argument for this formula is similar to equation (1). Putting equations 1-3 together, we get

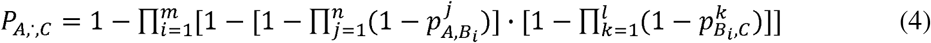

In Equation (4), *m* entities are l A and C, *n* instances of A-B_i_ relations in literature, and l instances of B_i_-C relations in literature. Extending this to multiple intermediate nodes between A and C is relatively straightforward. The above probabilistic framework will allow us to rank all the indirect relations that can be inferred. To infer the relation type (positive correlated or negative correlated) between two entities, which multiple intermediate entities could link, we use 1 to represent positive correlations, -1 to represent negative correlations, and 0 to represent unknown correlation type between any two entities connected by a direct edge and multiply all the correlations together. The resulting value, 1, -1, or 0 will give us the correlation type between the two entities. If there is at least one unknown correlation type (0) between the two entities, the overall correlation type is unknown. If there is no 0 and an even number of negative correlations, then the overall correlation type will be a positive correlation; otherwise, it is a negative one. For A-C entity pair to have a non-zero probability, there must be a path from A to C with all the directions going from A to C, such as A->B->D->C, while A->B<-D-=C is not a valid path from A to C.

In this manuscript, we show one application of PSR using our iKraph that calculates the indirect relationship between two entities: discover the repurposed drug. We present two study cases for identifying repurposed drugs for COVID-19 and cystic fibrosis, along with an additional study that involves predicting both repurposed drug candidates for 10 common diseases and potential additional uses for 10 common drugs. The details can be found in the Section 1.7 in SI. Fig. 5E illustrates Genistein’s repurposing for COVID-19 treatment, interacting with 25 human genes. It negatively affects 3 genes that have a positive impact on COVID-19 while positively influencing the remaining 22 genes, which are negatively associated with the disease. This evidence supports Genistein’s potential as a COVID-19 treatment candidate.

### 3. Integrating relations from public databases

To integrate the relations in the public databases, we downloaded the relations from two databases that have integrated data from a large number of databases recently, Hetionet^73^ and primeKG^92^, where Hetionet has integrated data from 29 databases and primeKG has integrated data from 20 databases. The total number of unique databases from both sources is 38. In addition, we extracted drug-target relations from the Therapeutic Target Database (TTD)^93^ and GO annotation^94,95^. In total, we integrated relation data from 40 public databases. The KG covers twelve common entity types: diseases, chemical compounds, species, genes/proteins, mutations, cell lines, anatomy, biological processes, cellular components, molecular function, pathway, and pharmacologic class. It covers fifty-three different relation types.

Among them, eight were annotated in the LitCoin dataset: association, positive correlation, negative correlation, bind, cotreatment, comparison, drug interaction, and conversion. Other relation types came from public databases. When incorporating relations from public databases to maintain the quality of the resulting KG, we excluded relations generated by high-throughput experiments, which are well-known to have a high proportion of false positives, and those predicted by previous machine learning models.

### 4. Incorporating relations from analyzing RNASeq data

We downloaded more than 300,000 human RNASeq profiles from the recount3 database^96^. We performed two types of analysis: differential gene expression analysis (DGEA) and gene regulatory network inference (GRNI). DGEA gave 92,628 differentially expressed genes for 36 different diseases, which correspond to 92,628 disease-gene relations, either positive or negative correlation, depending on whether the genes are up or downregulated in the corresponding diseases. GRNI gave 101,392 gene regulatory relations overall. We added close to 200,000 additional relations by analyzing this RNASeq dataset.

## Data Availability

The datasets used in this study are available on the GitHub repository at https://github.com/myinsilicom/iKraph^97^. Due to size limitations, additional large datasets can be accessed via Zenodo at https://zenodo.org/records/14851275^98^. We used BioRED dataset to train our NER and RE models, and the BioRED dataset can be accessed through https://ftp.ncbi.nlm.nih.gov/pub/lu/BioRED/. The complete knowledge graph is hosted on the cloud-based platform: https://www.biokde.com. The downloadable version of the complete iKraph can be accessed via Zenodo at https://zenodo.org/records/14851275^98^.

## Code Availability

The code and datasets generated during this study can be found via the GitHub repository at: https://github.com/myinsilicom/iKraph^97^.

## Acknowledgments

We thank the LitCoin NLP Challenge organizers for generating the valuable challenge dataset, which made the work possible.

This research was partially supported by the National Institutes of Health (NIH) under grant R21LM014277 (JZ), contract 75N91024C00007 (JZ), and contract 75N93024C00034 (JZ); by the National Science Foundation (NSF) under grants 2335357 (JZ) and 2403911 (JZ); and by the National Cancer Institute, National Institutes of Health, under Prime Contract No. 75N91019D00024, Task Order No. 75N91024F00030 (JZ). The content of this publication does not necessarily reflect the views or policies of the Department of Health and Human Services, nor does mention of trade names, commercial products, or organizations imply endorsement by the U.S. Government. The funders had no role in the study design, data collection and analysis, decision to publish, or preparation of the manuscript.

## Author Contributions Statement

Y.Z., X.S., F.P., and K.Y. contributed equally to this work. Y.Z., X.S., F.P., K.L., S.T., A.E., Q.H., W.W., Jianan W., and Jian W. collected data and developed models and pipelines. Y.Z., F.P., and J.Z. analyzed the data and developed methods. D.S., H.C., J.Zh., E.Z., B.L., T.Z., and J.Z. developed the iExplore platform interface. K.Y. and J.Z. conceptualized and designed the study. Y.Z., F.P., K.Y., and J.Z. wrote the manuscript. X.Q., T.Z., and P.Z. provided consultation and manuscript revision. J.Z. supervised the study and is the corresponding author.

