## Supplementary material for "A comprehensive large scale biomedical knowledge graph for AI powered data driven biomedical research": SI

### 1. Methods

The construction of iKraph involves four primary stages: named entity recognition (NER), entity normalization, relation extraction (RE), and novelty classification. Our approach employed fine-tuning pre-trained language models such as RoBERTa[1] and pubmedBERT[2]. Both single-sentence and multi-sentence modeling strategies were utilized in the RE task. The procedure of our method is depicted in Fig. S1.

#### 1.1 Named Entity Recognition (NER)

Each abstract was divided into individual sentences. For the annotation of training sentences, a BIO (Beginning, Inside, Outside) tagging scheme was used to label each word. Along with the BIO tags, each entity received annotations indicating its type (detailed in Table S1). This information was subsequently fed into RoBERTa for optimization, ensuring that the model assimilated the most pertinent data from our samples.

Next, the fine-tuned model was employed to predict the type-B/I/O tagging for each word. To obtain information at different levels, various combinations of hyper-parameters were set for RoBERTa, and prediction results were generated for each instance. The results were then consolidated using a majority vote.

Sometimes, the predicted results could be inconsistent, producing technically impossible sequences like "B I O I." To rectify such cases, a post-processing step was implemented. Rule-based label refinement and corrections were applied to ensure that the results did not breach linguistic guidelines, such as the requirement for a sequence of 'I's to follow a 'B', 'O' should not be between 'B' and 'I', etc. Ultimately, sequences starting with 'B' followed by either nothing or a sequence of 'I's were extracted as predicted entities.

#### 1.2 Entity Normalization

In the LitCoin challenge, the IDs of the entities were given, so there was no need for entity normalization (mapping all the synonyms of an entity to a unique ID). To construct a general KG for all PubMed abstracts, we need to map all the tagged entities to unique IDs. In BioCreative Challenge VII, our method ranked second for chemical term normalization and ranked first overall for chemical NER, normalization, and indexing[3,4].

To apply our NER pipeline to all PubMed abstracts, we obtained 34,531,488 documents with titles and abstracts from the PubMed database. The latest documents included were published in May 2023. Using our NER model, six types of entities were tagged in the tokenized texts: diseases, genes/proteins, chemical compounds, species, genetic variants, and cell lines. The goal of entity normalization is to assign proper entity IDs to the tagged entities to build a database of all entities, which should be consistent with existing databases and perform well in mapping synonyms of an entity to the same ID.

##### 1.2.1 Handling abbreviations

Abbreviations are widely used in scientific literature. However, the same abbreviation could represent different terms in different articles. For example, AS can refer to two different diseases, Ankylosing Spondylitis or Asperger Syndrome, in different articles. Correctly mapping abbreviations to their full names is critical for assigning correct entity IDs. To get the full names of abbreviations, we used Abbreviation Plus Pseudo-Precision (Ab3P)[5], which has a precision of over 0.8. We usually only map abbreviations to full names when they are standalone words. However, it is common for abbreviations to have a plural form; in such cases, we map the plural form of an abbreviation to the same entity ID. For example, if AD refers to Alzheimer's Disease, ADs will be expanded as Alzheimer's Diseases and mapped to the same ID of Alzheimer’s Disease.

1.2.2 Handling capitalizations

We map capitalized words and non-capitalized words to the same ID except for gene names and cell lines, which are treated differently. For example, by gene naming convention, TP53, Tp53, and tp53 are the same gene from different species with different IDs. As a result, the exact forms of gene names were used to identify their IDs. In fact, TP53 itself can have multiple IDs for different species. Handling of this is described later in section “Assign entity ID in NER results”.

1.2.3 Synonym dictionaries

To map synonyms to their correct IDs, for each ID, we need to identify as many synonyms as possible. We used data from multiple sources and evaluated the quality and quantities of each data source to create a priority list to handle conflicts. The following databases were used: MeSH[6], PubTator[7], MONDO[8], DOID[9], UMLS[10], SYMP[11], HP[12], OMIM[13], PubChem[14], DrugBank[15], Hetionet[16], PrimeKG[17], and Cellosaurus[18]. The priority list for each entity type is given below, where A > B means we will use A first and then B:

- Disease: MeSH (concept) > MONDO > DOID > UMLS > MONDO_grouped > SYMP > HP > OMIM > PubTator (MeSH (descriptor) > MeSH (supplement) > OMIM)
- Gene: PubTator (NCBI)
- Chemical: PubChem > MeSH (concept) > Drugbank > PubTator (MeSH (descriptor) > MeSH (supplement))
- Species: PubTator (NCBI Taxonomy)
- Genetic Variant : PubTator (tmVar[19])
- CellLine: Cellosaurus > PubTator (Cellosaurus)

1.2.4 Handling plural forms

We map the plural form of terms to their singular forms, if possible, using Python pattern package. We avoid abbreviated term when the ending “s” is capitalized (AIDS will not be considered as the plural form of AID).

After the above procedure, each set of synonyms in the dictionary is mapped to a unique ID (except genes).

#### 1.3 Assign entity ID in NER results

The process for assigning entity IDs to words tagged in the NER results is outlined below:

1. Verify if the word contains an abbreviation from the abbreviation list; if it does, expand the abbreviation as per the guidelines in the previous section.
2. Other than Gene and CellLine, convert capitalized terms to lowercase.
3. For any given term, we examine four distinct variations, prioritized as follows: the term's original form, its singular version, its lowercase rendition, and finally, a form that's both singular and lowercase. The search for these versions in the synonym dictionary begins with the highest priority form. The first version identified dictates the entity ID assignment. If the dictionary search yields no results, we utilize the transformed terms in seeking an ID. Should all versions be absent in the dictionary, we'll generate a new ID, adopting the original term as the entity name without regard to case distinctions.

The approach to identifying Gene entities is distinct due to their unique nature. Many terms within this category correspond to multiple IDs, reflecting genes across various species. To address this complexity, we implemented a series of refinement techniques, prioritized as follows:

- Identifying the closest Species mention within the same sentence to determine the target species. If there's an associated ID between the gene mention under review and the target species, this specific ID is selected.
- Employing the 'gene2pubmed' data file from the NCBI database, we cross-reference a compilation of gene IDs associated with a particular PubMed ID. If there's a match between any gene ID from this resource and those in our dictionary, that particular ID is chosen.
- Analyzing PubTator findings as a benchmark, if PubTator annotations also recognize a mention as a gene and associate it with a distinct ID, we then adopt this ID.
- If these strategies do not yield a result, we default to selecting the gene ID that appears most frequently within the PubTator annotations.

Upon completing these steps, each entity flagged in the NER process receives a definitive ID, setting the stage for subsequent relation extraction phases.

#### 1.4 Relation Extraction (RE)

The annotation of relations was primarily conducted at the sentence level, meaning that two entities were annotated with a potential relation only if they appeared in the same sentence. If a relation did exist, its type was identified. Sentence-level annotation was supplemented with document-level annotation, which did not require entities to be in the same sentence, thus allowing for more pairs of entities to be examined within a document.

The data annotated at the sentence level was processed through PubMedBERT for relation classification. This classification was bifurcated into two stages: firstly, determining the existence of a relation between a pair of entities and then identifying the specific type of relation. Since the method worked sentence by sentence, there could be discrepancies when a pair of entities occurred in multiple sentences. To reconcile such issues, a post-processing regression model was utilized to consider the predicted probability across all occurrences, deriving a final determination.

A multi-sentence model was also developed because not all related entities may appear in the same sentence. It gathered all sentences containing the paired entities and concatenated them into a single input for final prediction. It's important to note that the multi-sentence model's performance was 2-3% lower in the F1 score, so it was primarily utilized for pairs that never occurred in the same sentence and where the single-sentence model was not applicable.

In alignment with the NER task, these procedures were implemented with varied hyper-parameters to capture information as comprehensively as possible. A majority vote was then used to finalize the results, ensuring a robust and nuanced approach to relation annotation within the iKraph framework.

#### 1.5 Novelty Classification

The task of novelty classification is approached as a binary classification problem. It seeks to determine whether a triplet (consisting of a pair of entities along with their relation) is novel, as declared by the author of the article, opposed to the background knowledge. To accomplish this, all sentences from the same abstract containing the target pair of entities are concatenated together to form the training sample.

The samples are then processed through RoBERTa, which is fine-tuned for a binary classification task. To ensure a comprehensive assessment, multiple combinations of hyper-parameters are employed, capturing information at various levels.

The final determination of novelty is reached by employing a majority vote across the different models.

#### 1.6 LitCoin NLP Challenge performance

For NER, we ranked 2nd with a score of 0.9177. The first and third-ranked teams had scores of 0.9309 and 0.9068, respectively. For RE, we ranked 1st with a score of 0.6332 (modified Jaccard score). The second and third-ranked teams had scores of 0.5941 and 0.5681, respectively. NER counts for 30% of the total score, and RE counts for 70%.

#### 1.7 Case study: Discover repurposed drugs.

To pinpoint drugs potentially repurposed for treating disease A, our initial step was to pull all genes that exhibited either a positive or negative correlation with the disease. Within this scope, we primarily focused on relationships that were rooted in genes and targeted the disease. These associations could span multiple publications. For every publication, we calculated a relation probability, represented as pi. The cumulative probability and score for a connection between a gene and the disease are expressed as:

$$P_{gene \to disease} = 1 - \prod_{i=1}^{N} (1 - p_i) \tag{5}$$

$$S_{gene \to disease} = -\sum_{i=1}^{N} \log\,(1 - p_i + 0.01) \tag{6}$$

In cases where positive and negative correlations were present, the relationship's nature was determined by comparing the probabilities. If the positive correlation's probability surpasses that of the negative one, then the relationship is defined as positively correlated, and both the probability and score are set accordingly. Conversely, if the positive correlation's probability is lesser, the relationship is deemed negatively correlated, adjusting the probability and score in line with this finding. If probabilities for both correlations are equal, the score becomes the deciding factor.

After this, we indexed drugs that had correlations (be they positive or negative) with the genes we previously detailed. Ensuring the relationship's flow from the drug towards the gene, we noted that just like earlier, each drug-gene pair might be referenced in several publications, each assigned its unique relation probability, $p_i$. The overall probability and score for a link between a drug and a gene are depicted as:

$$P_{drug \to gene} = 1 - \prod_{i=1}^{N} (1 - p_i) \tag{7}$$

$$S_{drug \to gene} = -\sum_{i=1}^{N} \log\,(1 - p_i + 0.01) \tag{8}$$

We apply a threshold (*P_cutRE*) to the probability $P_{gene \to disease}$ and $P_{drug \to gene}$, retaining only values above this threshold and setting all others to 0, effectively disregarding those relationships. The criterion for a drug to be deemed repurposed hinges on the relationship between drug-gene and gene-disease being opposite, such as one being positive and the other, negative. The comprehensive probability and score for a drug's association with the disease through a specific gene, $G_j$, are:

$$P^{j}_{drug \to disease} = P^{j}_{drug \to gene} \cdot P^{j}_{gene \to disease} \tag{9}$$

$$S^{j}_{drug \to disease} = S^{j}_{drug \to gene} \cdot S^{j}_{gene \to disease} \tag{10}$$

Considering a scenario where the drug and disease are interconnected by M genes, the overarching probability and score are:

$$P_{drug\to disease} = 1 - \prod_{j=1}^{M}(1 - P^{j}_{drug\to disease}) \quad (11)$$

$$S_{drug\to disease} = \sum_{j=1}^{M} S^{j}_{drug\to disease} \quad (12)$$

Drugs that manifest an aggregate probability greater than 0.8 are identified as potential candidates for repurposing.

To determine if a candidate is already reported, we calculate the probability of drug-disease pairings, following the drug to disease direction. For each study, we determine a relation probability, denoted as pi. The total probability and the score linking a drug to a disease are defined as follows:

$$P_{durg\to disease} = 1 - \prod_{i=1}^{N}(1 - p_i)$$

$$S_{drug\to disease} = -\sum_{i=1}^{N} \log(1 - p_i + 0.01)$$

When assessing relationships with both positive and negative correlations, the nature is decided by comparing their probabilities. A relationship is considered negatively correlated if the probability of the negative correlation exceeds the positive one by 0.1, leading to categorizing the drug as reported for the disease. We apply a threshold (*P_cutDirect*) to the probability $P_{disease\to disease}$, retaining only values above this threshold and setting all others to 0, effectively disregarding those relationships.

##### 1.7.1 Drug repurposing prediction and validation

###### *1.7.1.1 Prediction (from Disease to find repurposed drugs)*

1. Input Parameters:
   Disease (D): **The target disease for drug repurposing.**
   Time Cutoff (YYYY-MM-DD): Specifies the cutoff date for including literature-based relations.
2. Relation Extraction:
   Retrieve all gene-disease correlations for disease D from iKraph, using only literature **published before the cutoff date**. These correlations are directed relations (gene->D) and can be:
      **Positive correlation**: The gene positively impacts the disease.
      **Negative correlation**: The gene negatively impacts the disease.
3. Drug Candidate Identification:
   For each gene identified:
      If the gene is positively correlated with D:
         Retrieve all drugs negatively correlated with the gene (drug->gene) as potential repurposed drug candidates.
      If the gene is negatively correlated with D:
         Retrieve all drugs positively correlated with the gene (drug->gene) as potential repurposed drug candidates.
4. Candidate Compilation:
   Compile a list of all drugs identified across all genes associated with D. These drugs are the predicted candidates for repurposing.

###### *Validation of Predictions*

1. Post-Cutoff Relation Extraction:

Using literature published after the cutoff date, retrieve all drug-disease correlations (drug->D) to validate the predicted candidates.

2. Validation Criteria:
   A drug is considered validated if it negatively correlates with disease D (indicating a treatment or potential therapeutic effect).

**Algorithm:** Drug Repurposing Prediction and Validation (from Disease)

```
procedure Drug_Repurposing(Disease, Time)
  PredictedDrugs ← ∅
  GeneDiseaseRelations ← GetGeneDiseaseRelations(Disease, Time)
  for each (gene, correlation) ∈ GeneDiseaseRelations do
      if correlation == "positive" then
          CandidateDrugs ← GetDrugGeneRelations(gene, "negative", Time)
      else if correlation == "negative" then
          CandidateDrugs ← GetDrugGeneRelations(gene, "positive", Time)
      end if
      PredictedDrugs ← PredictedDrugs ∪ CandidateDrugs
  end for
  PredictedDrugs ← RemoveDuplicates(PredictedDrugs)

  ValidatedDrugs ← ∅
  DrugDiseaseRelations ← GetDrugDiseaseRelations(Disease, Time, after_cutoff=True)
  for each drug ∈ PredictedDrugs do
      if drug ∈ DrugDiseaseRelations and DrugDiseaseRelations[drug] == "negative" then
          ValidatedDrugs ← ValidatedDrugs ∪ {drug}
      end if
  end for

  return PredictedDrugs, ValidatedDrugs
end procedure
```

###### *1.7.1.2 Prediction (from Drug to find repurposed indications)*

1. Input Parameters:
   Drug (X): **The target drug for repurposing.**
   Time Cutoff (YYYY-MM-DD): Specifies the cutoff date for including literature-based relations.
2. Relation Extraction:
   Retrieve all drug-gene correlations for drug X from iKraph, using only literature published before the cutoff date. These correlations are directed relations (drug->gene) and can be:
   - **Positive correlation**: The drug positively impacts the gene.
   - **Negative correlation**: The drug negatively impacts the gene.
3. Disease Candidate Identification:
   For each gene identified:
   - If X is positively correlated with the gene:

Retrieve all diseases negatively correlated with the gene (gene->disease) as potential indications.

If X is negatively correlated with the gene:

Retrieve all diseases positively correlated with the gene (gene->disease) as potential indications.

4. Candidate Compilation:
   Compile a list of all diseases identified across all genes associated with X. These diseases are the predicted repurposed indications.

***Validation of Predictions***

1. Post-Cutoff Relation Extraction:
   Using literature published after the cutoff date, retrieve all drug-disease correlations (drug->D) to validate the predicted candidates.
2. Validation Criteria:
   A drug is considered validated if it shows a negative correlation with disease D (indicating a treatment or potential therapeutic effect).

**Algorithm:** Drug Repurposing Prediction and Validation (from Drug)

```
procedure Drug_Repurposing(Drug, Time)
   PredictedIndications ← ∅
   DrugGeneRelations ← GetDrugGeneRelations(Drug, Time)
   for each (gene, correlation) ∈ DrugGeneRelations do
      if correlation == "positive" then
         CandidateDiseases ← GetGeneDiseaseRelations(gene, "positive", Time)
      else if correlation == "negative" then
         CandidateDiseases ← GetGeneDiseaseRelations(gene, "negative", Time)
      end if
      PredictedIndications ← PredictedIndications ∪ CandidateDiseases
   end for
   PredictedIndications ← RemoveDuplicates(PredictedIndications)

   ValidatedDrugs ← ∅
   DrugDiseaseRelations ← GetDrugDiseaseRelations(Drug, Time, after_cutoff=True)
   for each disease ∈ PredictedIndications do
      if disease ∈ DrugDiseaseRelations and DrugDiseaseRelations[disease] == "negative" then
         ValidatedIndications ← ValidatedIndications ∪ {disease}
      end if
   end for

   return PredictedIndications, ValidatedIndications
end procedure
```

##### 1.7.2 Drug repurposing for COVID-19

Since COVID-19 is a global medical emergency, urged on timely drug progress, the publication and discoveries for COVID-19 bloomed during the first few years and even during the first few months. To predict repurposed drug candidates for COVID-19, we use articles published before a chosen cutoff date, such as the first day of April, May, or June, to make predictions. Papers published after the cutoff date are then used to validate these predictions. Additionally, if a predicted drug candidate had already been reported in articles published before the cutoff date, it is excluded from the prediction results. The *P_cutRE* for the repurposing is set as 0.8. We then use all the publications before 04/01/2023 to get the reported drugs for COVID-19, which is used to validate the predictions. The probability for the cutoff, *P_cutDirect,* is set as 0.5. We also downloaded the drugs under (or ever under) clinical trials regardless the phases by 2023/07 for COVID-19 from https://clinicaltrials.gov/ to validate the predictions. We verified the drug candidates identified for repurposing by May 1, 2020, to see if they were subsequently reported from June 1, 2020, through July 1, 2023, monthly. 50% of candidates have been reported in the first 5 following months.

The last date of a month or a year was selected for monthly or yearly validation, and the publication date was extracted from the PubMed database for each abstract. Once a cutoff date is selected, all the publications before that date were used for drug repurposing prediction, and all the publications after that date were used for validation.

##### 1.7.3 Drug repurposing for cystic fibrosis

Yearly predictions for repurposed drug candidates targeting cystic fibrosis were conducted from 12/31/1985 to 12/31/ 2022, using PubMed publications available before each year's end as the basis for prediction. All the predictions reported before the cutoff time will be removed from the candidate. These candidates were then reviewed to determine if they had been reported post-cutoff. A consistent probability threshold of 0.5 was applied for both the phases of the prediction (P_cutRE) and validation (P_cutDirect).

##### 1.7.4 Drug repurposing for 10 common drugs and 10 common diseases

We predict the repurposed drug candidates for 10 common diseases and repurposed indications for 10 common drugs using all the publications by 12/31/2022. The result candidates contain the reported ones before the cutoff time. We reported the total repurposed and the reported drugs (indications). For each target disease or drug, we also calculate the total reported treatment or indications. The probability threshold is set as P_cutRE=0.8 and P_cutDirect=0.9.

We manually read a small number of predictions. There are two types of false negatives. First, the information extraction methods can make wrong predictions in any of the following areas: NER, RE, entity normalization, and direction prediction, which will generate false positives. Second, some of the repurposed drugs are quite toxic, indicating that toxicity should be considered either explicitly or implicitly in future studies.

#### 1.8 Integrating relations from public databases

To integrate the relations in the public databases, we downloaded the relations from two databases that have integrated data from a large number of databases recently, Hetionet[16] and primeKG[17], where Hetionet has integrated data from 29 databases and primeKG has integrated data from 20 databases. The total number of unique databases from both sources is 38, which including the following sources: Mayo Clinic, Orphanet[20], DisGeNET[21], human PPI network compiled by Menche *et al* [22], BioGrid[23], STRING[24], HuRI[25], Reactome[26], Bgee[27], Disease Ontology(DO)[28], MONDO[29], Entrez Gene[30], Gene Ontology[31], Human phenotype ontology (HPO) [12], Uberon[32], UMLS knowledgebase[10], Comparative Toxicogenomics Database (CTD)[33], DrugCentral[34], DrugBank[15], SIDER[35], MeSH, WikiPathways[36], LabeledIn[37], MEDLINE, Pathway Interaction Database[38], DISEASES[39], GWAS Catalog[40], TISSUES[41], BindingDB[42], MEDI[43], PREDICT[44], DOAF[45], EHRLink[46], Evolutionary Rate Covariation[47], Hetio-dag[48], Incomplete Interactome, Human Interactome Database[49–52], STARGEO[53]. In addition, we extracted drug-target relations from the Therapeutic Target Database (TTD)[54] and GO annotation[55,56]. In total, we integrated relation data from 40 public databases. The KG covers twelve common entity types: diseases, chemical compounds, species, genes/proteins, mutations, cell lines, anatomy, biological processes, cellular components, molecular function, pathway, and pharmacologic class. It covers fifty-three different relation types. Among them, eight were annotated in the LitCoin dataset: association, positive correlation, negative correlation, bind, cotreatment, comparison, drug interaction, and conversion. Other relation types came from public databases. When incorporating relations from public databases to maintain the quality of the resulting KG, we excluded relations generated by high-throughput experiments, which are well-known to have a high proportion of false positives, and those predicted by previous machine learning models.

### 2. Evaluation of the large-scale information extraction

**Human Annotators**:

Two Ph.D. annotators constructed the gold standard. They obtained their Ph.D. in physics focusing on biophysics from North Carolina State University and have since continued their career in the biomedical domain. The annotations with conflicts between the two annotators were then processed by a professor of biostatistics, who has been working on biomedical problems for over 15 years, to finalize the ground truth.

For our benchmark data, we selected a random sample of 50 PubMed abstracts, including 1583 entity pairs. Of these pairs, 151 could be identified and categorized within our established set of 8 relationship types, with 97 of these marking discoveries. The results shown in Table 1 were based on this ground truth, and the result of the experienced annotator was taken from the result used to construct the gold standard. E-merge Tech provided a commercial professional annotation service with a long history of servicing the annotation of biomedical text mining tasks.

### 3. Analyzing Network Structure in Our iKraph

Figure 1E is the degree distribution of our iKraph, wherein k represents the degree of an individual entity, and p(k) denotes the corresponding probability of any entity exhibiting that degree (Figure 1E). The structural topology of our network adheres to a power-law distribution, as indicated by an alpha parameter value of 3.003. A methodical comparative assessment with exponential distributions underscores the preeminence of the power-law model, substantiated by an R-value of 22.2196 and a remarkably significant p-value approximating 2.22e-9. These metrics convincingly dispel the notion of serendipitous network connectivity distribution, emphasizing instead a pronounced power-law characteristic.

To fortify our theoretical postulation with empirical evidence, we utilized the Kolmogorov-Smirnov statistical test. This rigorous comparison between our theoretical power-law distribution and collected empirical data yielded a K-S statistic of 0.0 and a p-value of 1.0, signifying an exceptional degree of agreement. These results compellingly reinforce the thesis that the network under investigation exhibits a scale-free topology, with its linkages rigorously conforming to a power-law distribution.

#### 4. The interface of iKraph

We have developed an interactive tool, called iExplore, at https://www.biokde.com for researchers to access iKraph with two main modules: direct and indirect relation search. A detailed user guide is provided on the website. Direct relations are those that are explicitly described in the literature, while indirect relations are inferred using our PSR algorithm. With direct relation search, researchers can input the first entity name (required), a second entity name (optional), and other optional conditions such as date range, relation type, novelty, and contexts. When a relation type can have different directions, users can also specify the direction type. Using context as filters allows researchers to select relations related to only certain context, such as species, diseases, or cell lines. Contexts of a relation are defined by whether a context term co-occurs with the relation in the same abstract. For indirect relation search, we have made inference for three types of entities important for drug discovery: genes/proteins, diseases, and chemical compounds (including drugs). Researchers can use this tool to find new targets for a disease or conduct drug repurposing for a disease or a chemical compound.

In addition to query functionality, we have built an annotation function for users to modify/correct the entities and/or relations in iKraph. It works as a crowdsourcing annotation tool, with an advantage that the participants are much more experienced and has more knowledge in the domain, so that they will be able to provide higher quality annotations.

#### 5. Drug repurposing for COVID-19 on iExplore interface

Using the iExplore tool at BioKDE, under the Indirect Relationship Search tab, users can search COVID-19 as a disease for Entity 1, Drug as the type of Entity 2, Negative Correlation as the Relationship Type, and direction from Entity 2 to Entity 1. One unique feature of PSR is it identifies explicitly the genes that connect COVID-19, and the potential drugs, where either the gene is positively correlated to COVID-19 with direction from gene to COVID-19 (the gene is a causal factor for COVID-19) and the drug is negatively correlated to the gene with direction from drug to the gene, or the gene is negatively correlation to COVID-19 with direction from gene to COVID-19 and the drug is positively correlated to the gene with direction from drug to the gene. In both cases, the inferred relation between the drug and COVID-19 will be negatively correlated (meaning a therapeutic effect), and the direction is from drug to COVID-19 (a causal relation from drug to COVID-19). Figure S2A shows the top seven drug candidates for a drug repurposing search for COVID-19 using the indirect relation search at iExplore. For all seven candidates, we have found hypothesis articles proposing that they can be used as potential treatments for COVID-19 (APC[57], Genistein[58], LY294002[59], AAT[60], Rosiglitazone[61], PD98059[62], and ghrelin[63]). The number of genes connecting these top seven candidates to COVID-19 ranges from 45 to 81. By default, only a small number of top genes will be shown for each candidate. Users can find all the genes by performing another indirect search by specifying the drug name. Figure 2B shows the genes (with a probability greater than 0.9) connecting COVID-19 and Genistein, one of the top candidates. Clicking an edge will display all the literature evidence of the corresponding relation so that a manual verification can be conveniently performed. In this search, we have filtered out all the drugs used or tested for treating COVID-19 as reported previously in PubMed literature. Therefore, the result consists of drugs that have not been tested for COVID-19, which can be potential novel discoveries. Being able to select only candidates which have not been used as treatments previously is another unique feature of our approach, as it requires a comprehensive knowledge of existing literature, which has yet to be possible in previous studies.

#### 6. Future work and challenges

There are several areas we can work on in the future to further enhance the coverage and functionalities of iKraph. Firstly, we extracted the relations from the abstracts of articles that are indexed in PubMed. The rationale is that the more important part of our biomedical knowledge was likely mentioned at least once in the PubMed abstracts. In addition, the models we developed for extracting relations from PubMed abstracts may also need to be further tuned to extract relations from full-text articles. As a large number of full-text articles are available from PubMed Central database, we plan to extract relations from this valuable resource in the near future, which will add a substantial amount of knowledge to iKraph; Second, we can add more analysis results from publicly available genomics datasets, such as those generated from NIH common fund initiatives; Third, although LitCoin dataset covers six major types of biological entities important for drug discovery and development, there are other key entity types that can be added that would enhance the capability of iKraph, such as organ/tissue names, biological pathways/processes, experimental methods, etc.. This would require additional efforts in annotating the entities and the new relation types introduced with the addition of the new entity types; Fourth, a substantial amount of our knowledge was deposited in supplementary materials of published articles, which are usually not indexed. Developing methods or a crowdsourcing mechanism to incorporate such knowledge will substantially increase the findability, accessibility, and reusability of such knowledge; Fifth, a crowdsourcing platform can be built to allow researchers to submit their new discoveries directly to iKraph, which will allow the timely access of new findings with higher quality as these are manually submitted. Such mechanism also allows submission of knowledge in supplementary materials; and finally, the PSR algorithm we designed for inferring indirect causal relations were only applied to certain types of entity pairs. Despite having produced many plausible hypotheses, it is also limited to predict relations for which the entities must be connected with specific directions. It also does not incorporate other rich information in other parts of the KG when making the inference. More general and more powerful methods, i.e. graph based deep learning methods, will likely take the application of iKraph to a new level.

We highlight some challenges we faced during this study to encourage the research community to address them to make knowledge graphs more powerful discovery tools. Firstly, although at instance level, our model has a very balanced precision and recall, the false positive rate can be dramatically inflated at entity pair level for large-scale information extraction. A pair of entities may co-occur in multiple PubMed abstracts. Each co-occurrence is called an instance of this entity pair. If any one of the co-occurrences is predicted as a certain relation, we will assign this relation to this entity pair even if all the other co-occurrences are predicted as no relations. For some common entities, even if there are no relations among them, they can co-occur in many different abstracts. Each occurrence will need to be predicted by the model and by chance some occurrences may be wrongly predicted, which will result in wrong relations between the entities. The PSR algorithm was designed to address this issue. But there is still a lot of room for further improvement. Another issue is entity normalization or entity linking, which maps synonyms of an entity to the same official name and ID. Currently, we are using high-quality databases and PubTator for entity normalization. However, from our manual checking of the extracted relations, substantial errors were associated with the noise introduced in the entity normalization step. More studies are needed to improve the performance of this step.

**Potential biases in KG construction.** The potential biases may come from the following areas: first, we did not extract information from full-text part of the published papers. There is information authors tend to describe in full-text articles, which will not be extracted out; Second, the models were trained using manually labeled data, which will be affected by the biases produced in the annotation guideline design and manual annotation process; Third, the information extracted from PubMed literature contains only 6 types of entities. Important entities and their relations such as biological processes, tissue/organ, cell types, etc. were only obtained from databases, which lack sufficient coverage of the full PubMed literature. These limitations will be addressed in our future studies.

#### 7. Figures

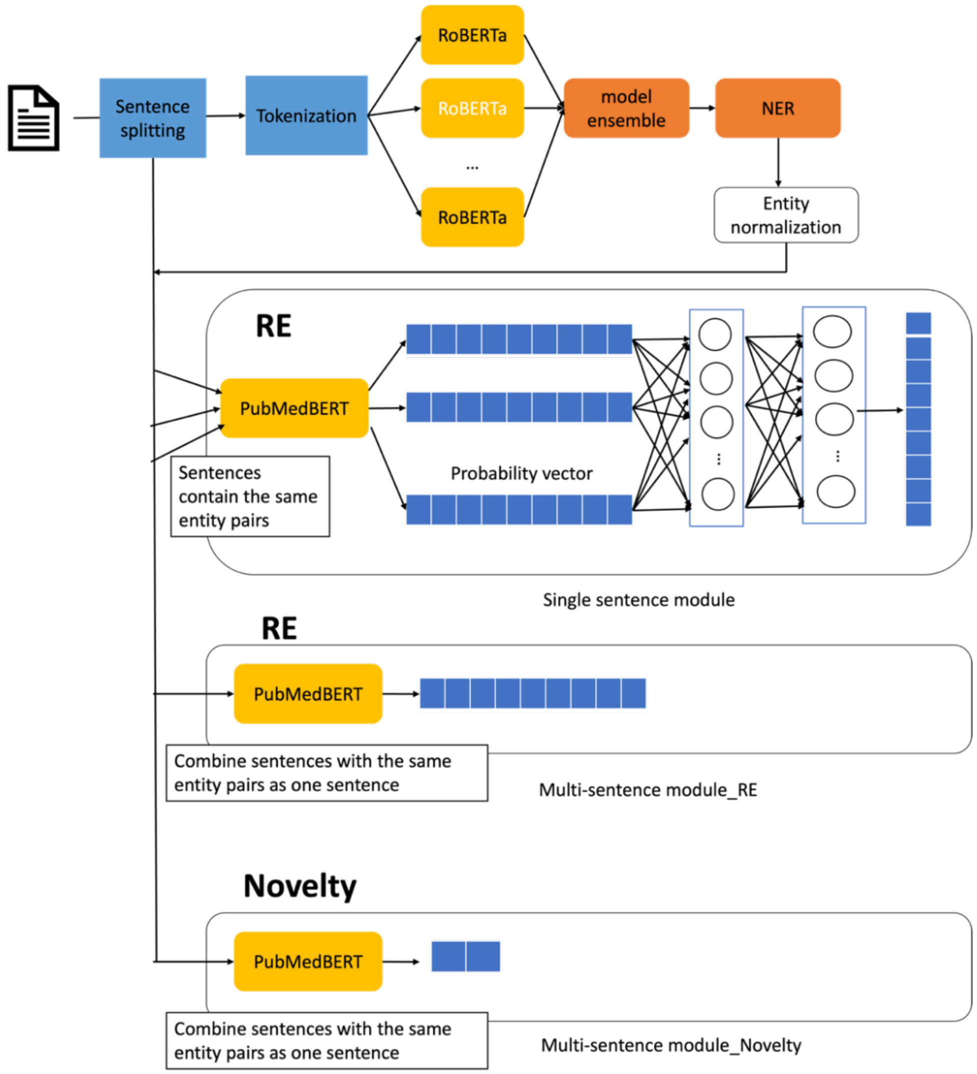

**Fig. S1**. The pipeline for knowledge graph construction.

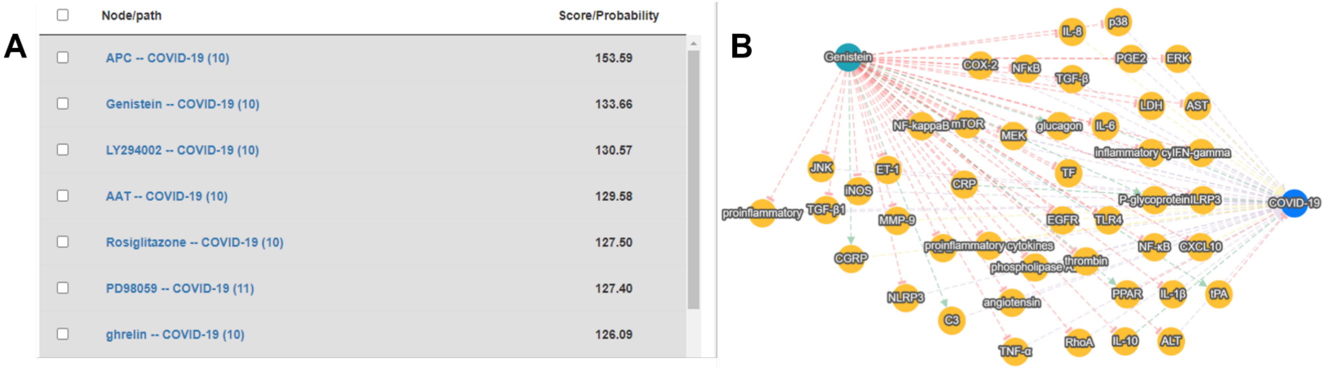

**Fig. S2. Drug repurposing using indirect relationship searches. A.** The result from searching COVID-19 as a disease for entity 1, drug as the type of entity 2, negative correlation as the relationship type, and entity 2 -> entity 1 as direction. By default, only a small number of top genes will be shown for each candidate drug/disease. We have found hypothesis articles for all seven candidates suggesting they can be used as potential treatments for COVID-19. **B.** Users can find more genes linking the drug and disease by performing another indirect search by specifying the drug name, for example, Genistein, in this case, which gives 81 genes connecting Genistein and COVID-19. The figure showed those with a probability greater than 0.9.

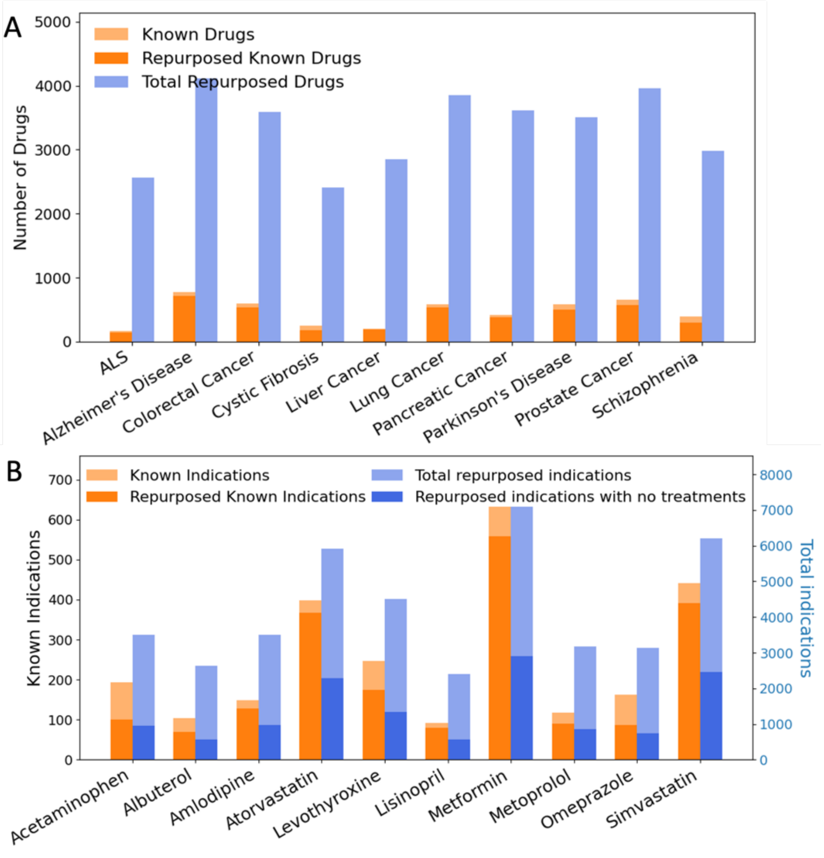

**Fig. S3**. **Drug repurposing for 10 diseases and 10 common drugs**. Our method identified large numbers of candidates. The known drugs or indications are those extracted from PubMed abstracts, not FDA-approved drugs or indications. On average, our method has repurposed more than 86% of the known drugs or 76% of indications. **A.** Drug repurposing for 10 diseases without satisfactory treatments; **B.** Drug repurposing for 10 common drugs.

Among the repurposed indications, some (dark blue bar) have not reported treatments in PubMed abstracts, which suggests unmet medical needs.

### 8. Tables

**Table S1**. A summary table of the types of concepts for the annotated entities.

| Entity type | Description | Example |
|---|---|---|
| Disease | Diseases, symptoms, and some disease-related phenotypes | Alzheimer Disease |
| Gene/Protein | Genes, proteins, mRNA, and other gene products | BRCA1(gene), Hemoglobin (protein) |
| Chemical Compound | Chemical compound and drugs | Donepezil hydrochloride, Glucose |
| Species | Species in the hierarchical taxonomy of organisms | Arabidopsis thaliana |
| Genetic Variant | Variations in the genome or proteins, including substitutions, deletions, insertions, and more | rs12979860 |
| CellLine | Cell line | MCF7/AdrR |

**Table S2: Overview of 11 hypothesized Treatments for COVID-19 Alongside Supporting Evidence**

| Drug Name | Evidence type | PubMed ID or ClinicalTrials.gov ID (starts with NCT) |
|---|---|---|
| PD98059 | Hypothesized in article | 34768113[64] |
| Vitamin D | Clinical trial | NCT04536298 |
| Rolipram | Hypothesized in article | 33623736[65] |
| Rosiglitazone | Hypothesized in article | 32344313[61] |
| Tacrolimus | Clinical trial | NCT04341038 |
| Genistein | Clinical trial | NCT04482595 |
| LY294002 | Hypothesized in article | 34345206, 34239361[59,66] |
| Vitamin E | Clinical trial | NCT04570254 |
| Alcohol | Clinical trial | NCT04719208 |
| U0126 | Hypothesized in article | 33011728[67] |
| DHA | Clinical trial | NCT04553705 |

**Table S3.** Comparative Performance of Model vs. Human Annotators on 50 PubMed Abstracts, encompassing 1583 entity pairs. Of these, 151 pairs were classifiable into one of our 8 relationship categories, with 97 out of the 151 representing novel discoveries. *Commercial annotation involved a subset of 10 PubMed abstracts randomly selected from the LitCoin Competition's training dataset. The P-value was calculated using the Wilcoxon signed-rank Test (two-sided).

| | RE task | Model | Experienced Ph.D. annotator | p-value (Model vs Ph.D.) | Commercial Annotation* |
|---|---|---|---|---|---|
| Relation | F1 | **0.7718** | 0.6568 | 0.0798 | 0.5385 |
| | Recall | **0.7616** | 0.7351 | 0.4010 | 0.6222 |
| | Precision | **0.7823** | 0.5936 | 0.0532 | 0.4746 |
| | Accuracy | **0.9629** | 0.9316 | 0.1454 | 0.8506 |
| Novelty | F1 | 0.9348 | **0.953** | 0.4428 | 0.7097 |
| | Recall | **0.9885** | 0.9342 | 0.2596 | 0.7097 |
| | Precision | 0.8866 | **0.9726** | 0.8015 | 0.7097 |
| | Accuracy | 0.904 | **0.9417** | 0.9133 | 0.6 |

**Table S4. Comparison of numbers of relations extracted from PubMed abstracts with those from public databases and co-occurrence. * Protein-protein interactions obtained using high-throughput experiments were not included since they are known to be very noisy.**

| Relation Type | public databases | iKraph | co-occurrence |
|---|---|---|---|
| Chemical - Gene | 111,222 | **5,725,179** | 46,910,606 |
| Chemical - Chemical | 1,337,757 | **4,246,231** | 114,473,578 |
| Gene - Gene | 795,601* | **3,940,720** | 84,987,749 |
| Chemical - Disease | 275,556 | **4,254,418** | 38,462,777 |
| Disease - Gene | 119,091 | **3,959,355** | 37,607,758 |

#### 9. Boxes

**Box S1**. The key advances in this study.

**Key advances powered and unique capabilities enabled by iKraph:**

- A large-scale, human-level information extraction on all PubMed abstracts. The pipeline won first-place in the LitCoin NLP challenge.
- Extracted large number of relations that were not documented in the public databases.
- Constructing a causal knowledge graph by predicting the direction (causality) of relations.
- Extraction of novel discoveries in published articles. Distinguishing novel discoveries and background knowledge enable applications that requires novelty-specific information, such as training automated knowledge discovery models.
- An interpretable probabilistic-based causal relation inference algorithm.
- Drug repurposing and drug target identification tools that discovered large numbers of plausible candidates with sufficient literature evidence.
- Both drug repurposing and drug target identification have satisfactory recall and precision, which have been difficult to estimate in previous studies due to incomplete retrieval of existing literature.
- Incorporating analysis results from high-throughput genomics data allows integrative analysis maximizing the available information in both literature and public experimental data.
- Demonstrated an effective approach for constructing databases from biomedical literature.
